# Transcriptional analysis identifies overlapping and tissue-distinct profiles between Kaposi sarcoma tumors of the skin and gastrointestinal tract

**DOI:** 10.1101/2022.03.18.484923

**Authors:** Ramya Ramaswami, Takanobu Tagawa, Guruswamy Mahesh, Anna Serquina, Vishal Koparde, Kathryn Lurain, Sarah Dremel, Xiaofan Li, Ameera Mungale, Alex Beran, Zoe Weaver Ohler, Laura Bassel, Andrew Warner, Ralph Mangusan, Anaida Widell, Irene Ekwede, Laurie T. Krug, Thomas S. Uldrick, Robert Yarchoan, Joseph M. Ziegelbauer

## Abstract

Kaposi sarcoma (KS), caused by Kaposi sarcoma herpesvirus (KSHV), is a multicentric tumor characterized by abnormal vasculature and proliferation of KSHV-infected spindle cells. KS commonly involves the skin but in severe cases KS can also involve the gastrointestinal tract (GI). Here, we sought to compare the cellular and KSHV gene expression signatures of skin and GI KS lesions. Skin and GI KS were compared to normal matched samples using bulk RNA sequencing.Twenty-two paired samples of KS and normal tissue were obtained (skin (10 pairs) and GI (12 pairs)) from 19 patients with KS of whom 17 had concurrent HIV infection. Seven paired samples were from patients who had received prior KS therapy. Three patients provided both skin and GI samples at the same timepoint. These analyses identified 370 differentially expressed genes unique to cutaneous KS and 58 DEGs unique to GI KS compared to normal skin or GI tissues. Twenty-six differentially expressed genes overlapped between skin and GI KS, which included *FLT4*, which encodes for a VEGF-C and VEGF-D receptor, and *STC1*. KSHV infection of primary lymphatic endothelial cells (LECs) resulted in increased angiogenesis, and repression of *STC1* or *FLT4* inhibited angiogenesis. The analyses of KSHV expression from KS lesions identified certain lytic genes, specifically ORF75, that were consistently expressed, and these expression patterns differed from laboratory infection of LECs with KSHV and KSHV gene expression in PEL cell lines. This study demonstrates that complex patterns of gene expression are found in KS tissue that differ from the canonical latent/lytic programs seen in KSHV cell lines and also demonstrates differences in viral gene and clinically relevant host gene expression in skin and GI KS that may offer insights into the pathogenesis of these forms of KS.

**One sentence summary:** Kaposi sarcoma that manifests in the skin and gastrointestinal tracts differ by human and viral gene expression.

## Introduction

Kaposi sarcoma (KS) is caused by Kaposi sarcoma herpesvirus (KSHV, also known as human herpesvirus 8 (HHV8)(1). In addition to KS, KSHV is the causative agent of primary effusion lymphoma (PEL), a form of multicentric Castleman disease (MCD), and the more recently identified KSHV inflammatory cytokine syndrome (KICS) (2–5). These conditions can occur alone or concurrently in the same patient and have different prognoses (6, 7). Within the United States, KS is seen predominantly among people living with HIV (PLWH), although the incidence of HIV-associated KS has decreased over time with advances in HIV treatment(8). However, KS remains one of the most common HIV-associated tumors in the US (noted as epidemic KS) and is one of the most common cancers overall in parts of sub-Saharan Africa, where endemic KS and epidemic KS contribute significantly to morbidity and mortality in this region (9).

Like other herpesviruses, KSHV generally exists either in a state of latency, in which a few of the viral genes are expressed and no infectious progeny are released, or lytic replication where most viral genes are expressed and infectious virions are produced and disseminate within the host(10, 11). Previous studies have shown that most of the spindle cells in KS lesions contain KSHV in its latent form, while a smaller number of cells express some KSHV lytic gene products (12, 13). KS lesions are notable for the proliferation of pathognomonic abnormal spindle cells, mixed inflammatory infiltrates, and the formation of aberrant, leaky vascular channels (14, 15). The spindle cells within KS lesions are infected with KSHV, and express vascular epithelial growth factor (VEGF) receptors and cytokines such as tumor necrosis factor-a and interleukin (IL)-6 (15–19). KSHV, like other herpesviruses, is notable for its molecular piracy of genes homologous to cellular regulatory genes (20, 21) and its modulation of cellular survival, angiogenic and immune regulatory pathways (22, 23).

KS most often occurs in PLWH where it is seen at low CD4^+^ T cell counts, or in the absence of HIV/AIDS, other immune dysregulation may contribute to onset of KS. KS most commonly manifests as skin lesions but may also occur in visceral organs including the respiratory and gastrointestinal (GI) tracts in severe cases(24, 25). Patients with GI KS often have gastrointestinal bleeding and anemia, as well as systemic symptoms and weight loss. The presence of KS in the GI tract indicate a diagnosis of a concurrent KSHV-associated disease such as KICS, PEL or MCD. These conditions are associated with elevated circulating IL-6, which can result in systemic inflammation and multiorgan dysfunction(3, 26, 27). The aim of KS treatment is disease control; treatment is usually administered until a plateau in response or until full resolution of KS lesions is achieved. Among PLWH, antiretroviral therapy (ART) is always a part of treatment (28). For more extensive disease or disease that does not respond to ART, systemic therapy is generally administered; this consists of chemotherapy, such as liposomal doxorubicin or paclitaxel, or more recently, pomalidomide, which is an immunomodulatory agent (29–31). Despite remission with systemic therapy, KS may recur, even in the presence of well-controlled HIV and higher CD4^+^ T-cell counts and require long-term, intermittent treatment. The reasons for recurrence in these instances are unclear. The lack of patient-derived KS cell lines and an animal model for KS has limited the ability to fully understand KS pathogenesis. The relationship between inflammation, immunity and the genomics of this disease has been examined in KSHV-infected lymphoma cell lines but not well-studied in KS tissues from patients with these disorders.

Little is known about cellular and viral RNA transcripts in KS lesions and how these factors are influenced by prior therapy, concurrent KSHV-associated diseases, or the tissue location of KS lesions. Recent reports used high throughput RNA sequencing to measure gene expression changes in KS in patients from sub-Saharan Africa. One study of both normal and KS tissue samples from four volunteers identified gene expression changes associated with glucose and lipid metabolism (32). Another study examined KS tumors in ART-naïve volunteers in Uganda(33). In this study of 23 cutaneous KS tumors, the authors reported that RNA from a single KSHV promoter within the latency region was highly expressed in the majority of KSHV transcripts from KS lesions.

We sought to understand how human gene expression and viral transcripts are altered in KS as compared to normal tissue using matched patient samples. In this study of participants with well-annotated clinical characteristics, we also investigated both skin and GI KS to determine differences in the viral and host transcripts between these lesions. We determined whether there were differences in immune environment by the location of the KS lesion or clinical characteristics. Specific cellular genes of interest that were linked to pathogenesis of KS or KSHV infection were investigated with *in vitro* studies using lymphatic endothelial cells (LECs) infected with KSHV to study and determine the significance of these findings from patient samples.

## Results

### Participant HIV and KS characteristics

Nineteen participants were included in this analysis, contributing 22 KS samples with paired normal tissue. There were 10 skin KS samples collected and 12 GI KS samples obtained from all participants (Figure 1A). Three participants provided both cutaneous and GI KS samples collected at the same timepoint and were included in these analyses. Seventeen participants (89%) had HIV-coinfection, 16 of whom were receiving antiretroviral therapy and at sample collection had a median CD4^+^ T cell count of 38 cells/μL and HIV viral load (VL) of 443 copies/mL at the time of sample collection (Table 1). In addition to KS, 10 participants met criteria for KICS (these patients did not have a diagnosis of PEL or MCD, diagnostic criteria noted in Table S1), 2 participants had concurrent MCD, 2 participants had PEL, and 2 participants had concurrent PEL and MCD. The median KSHV VL for all participants was 376.5 copies/10^6^ PBMCs. The median KSHV VL was 1148 copies/10^6^ PBMCs for those with GI KS and was 90 copies/10^6^ PBMCs for those with skin KS. Those with skin KS had a median HIV VL of 74 copies/mL and CD4^+^ T cell count of 38 cells/μL. Participants contributing GI KS lesions had a median HIV VL of 5134 copies/mL, CD4^+^ T cell count of 36 cells/μL.

**Figure 1:**
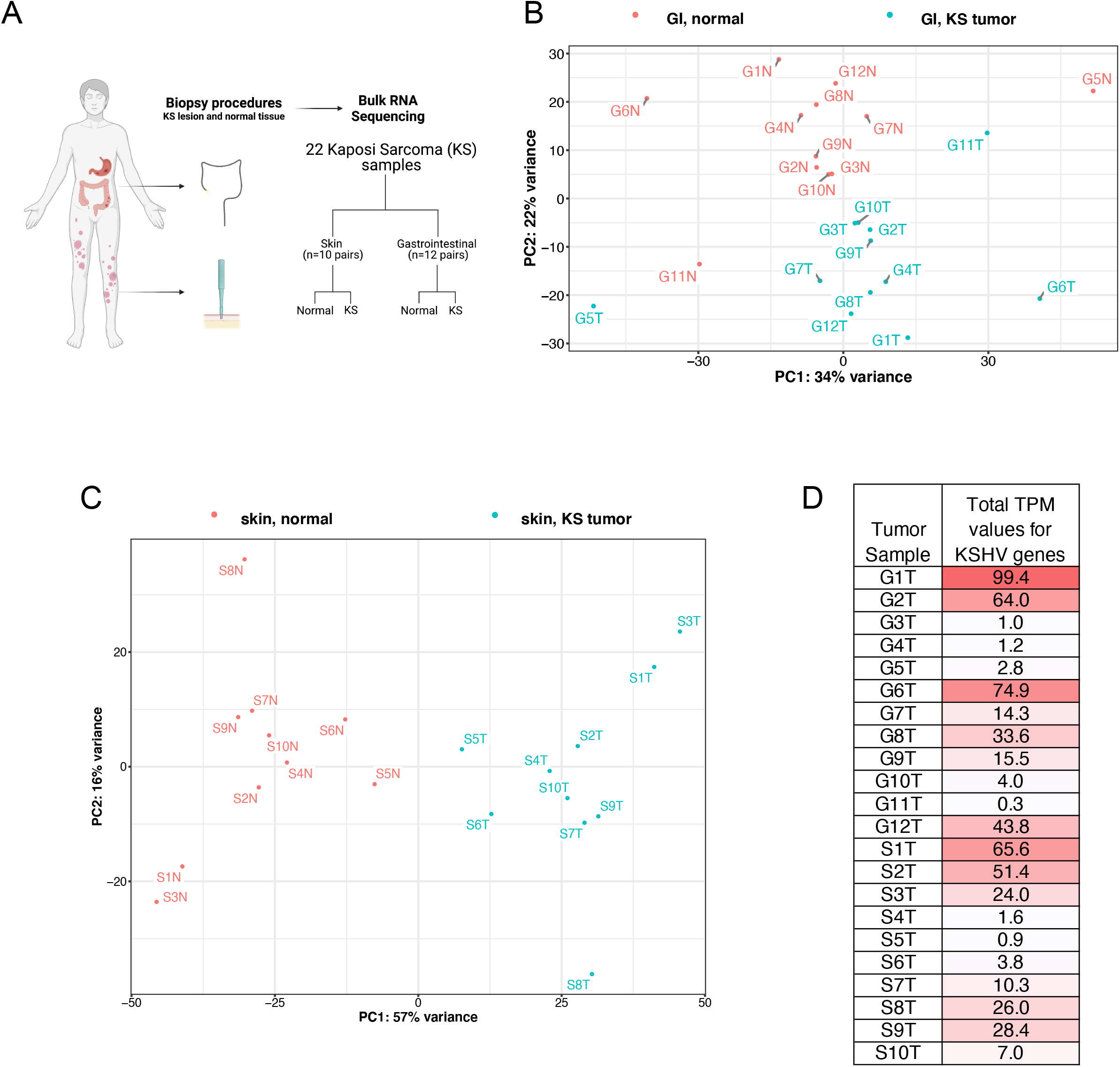
Sample types and principal component analysis of RNA expression profiles. A. Diagram of the normal and KS matched samples used in this study. After RNA sequencing, gene expression profiles of combined human and viral genes were analyzed using principal component analysis for skin samples (B) and GI samples (C). “N” at the end of sample names are the normal tissue samples. “T” in sample names refers KS tumor samples. D. KSHV transcript per million (TPM) values were combined for all KSHV genes. Stronger red backgrounds indicate higher KSHV expression.

**Table 1:** Baseline characteristics of all participants at the time of biopsy collection and characteristics by skin and GI sample collection. *Includes 3 participants who provided both skin and GI samples.

|  | **All participants**<br>**(N=19)** | **Skin KS participants\***<br>**(N=10)** | **GI KS participants\***<br>**(N=12)** |
| --- | --- | --- | --- |
| **Age** (median in years, IQR) | 37 (29, 46) | 43 (34, 49) | 30 (28.3, 36.1) |
| **Sex** (n, %) |  |  |  |
| Cisgender Male | 17 (89) | 10 (100) | 10 |
| Cisgender Female | 1 (5) |  | 1 |
| Transgender Female | 1 (5) |  | 1 |
| **Race** (n, %) |  |  |  |
| White | 4 (21) | 2 (20) | 3 |
| Black | 7 (37) | 3 (30) | 5 |
| Hispanic | 5 (26) | 4 (40) | 2 |
| African Immigrant | 3 (16) | 1 (10) | 2 |
| **HIV characteristics** |  |  |  |
| HIV co-infection (n, %) | 17 (89) | 9 (90) | 11 |
| Duration of HIV, median, (median in months, IQR) | 95 (5, 147) | 117 (5, 145) | 97 (26, 107) |
| CD4 T-cell count, cells/μL (median, IQR) | 38 (24, 100) | 39 (19, 195) | 36 (25, 85) |
| HIV viral load, copies/mL (median, IQR) | 443 (57, 5134) | 74 (39, 104) | 5134 (552, 47759) |
| On antiretroviral therapy (ART) at time of biopsy (n, % of HIV+) | 16 (94) | 10 (100) | 11 (92) |
| **KS characteristics** |  |  |  |
| Duration of KS diagnosis (median in months, IQR) | 5 (0.9, 32) | 5 (1, 38) | 10 (1, 16) |
| Prior therapy for KS (n, %) | 7 (37) | 3 (30) | 5 (42) |
| **Concurrent KSHV-associated diseases (n, %)** |  |  |  |
| None | 3 (16) | 3 (30) | 1 (8) |
| KICS | 10 (53) | 5 (50) | 7 (58) |
| MCD | 2 (11) | 1 (10) | 1 (8) |
| PEL | 2 (11) | 1 (10) | 1 (8) |
| MCD and PEL | 2 (11) | - | 2 (17) |
| KSHV-VL, copies/10<sup>6</sup> PBMCs (median, IQR) | 377 (0, 1798) | 90 (0, 728) | 1148 (0, 2792) |

### Differences and overlap in differentially expressed genes in GI and skin KS lesions in compared with paired normal samples

For all patients samples, total RNA was purified and RNA sequencing was performed. First, gene expression patterns were examined in individual samples using principal component analysis (PCA), which revealed that samples were distinguished by whether they came from KS lesions or normal tissue (Fig. 1B-C). To measure overall levels of KSHV gene expression for each lesion, total transcript per million (TPM) values were calculated (Fig. 1D). There was over a hundred-fold variation in the amount of KSHV transcripts detected in KS lesions in these samples. The large range in KSHV expression is likely attributable to varible levels of infected cells per tissue overall, and then different proportions of latent or lytically-infected cells. Overall, KS lesions with higher numbers of KSHV transcripts (Fig. 1D) displayed a larger separation between matched tumor and normal samples in the PCA plots (Fig. 1B-C), reflecting a larger difference in overall gene expression patterns.

To determine the gene expression difference between each KS sample and the matched normal tissue sample from the same participant, cellular genes that had a greater than log2 fold change of 2.0 with an adjusted p-value less than 0.05 were determined (Figure 2). Of approximately 17,000 mapped genes, there were 370 differentially expressed genes (DEGs) unique to skin KS and 58 DEGs unique to GI KS, and 26 common to both as compared to their respective normal tissues (Figure 2A). A pathway enrichment analysis was performed on the lists of DEGs in skin and GI samples (Table S2-S3). One of the most enriched pathways in both set of samples included the granulocyte adhesion and diapedesis pathway. IL-6 signaling and HIF1a signaling pathways were also enriched in both skin and GI KS samples. The B cell receptor signaling pathway was enriched in the skin KS samples, but not the GI KS samples. Of note, many enriched pathways contained IL1A and various matrix metallopeptidases. The 26 DEGs that were shared in both the GI and skin samples as compared to their matched normal samples (Fig. 2A-B). Only one gene, *OTOP3*, was repressed within KS lesions as compared to the normal counterparts. A striking difference between skin and GI KS was that that IL1A was increased in GI KS but repressed in cutaneous KS (Fig. 2B). Additional analyses identified a correlation between IL1A expression and overall levels of KSHV expression in GI KS, which was not observed in skin samples (Table S4).

**Figure 2:**
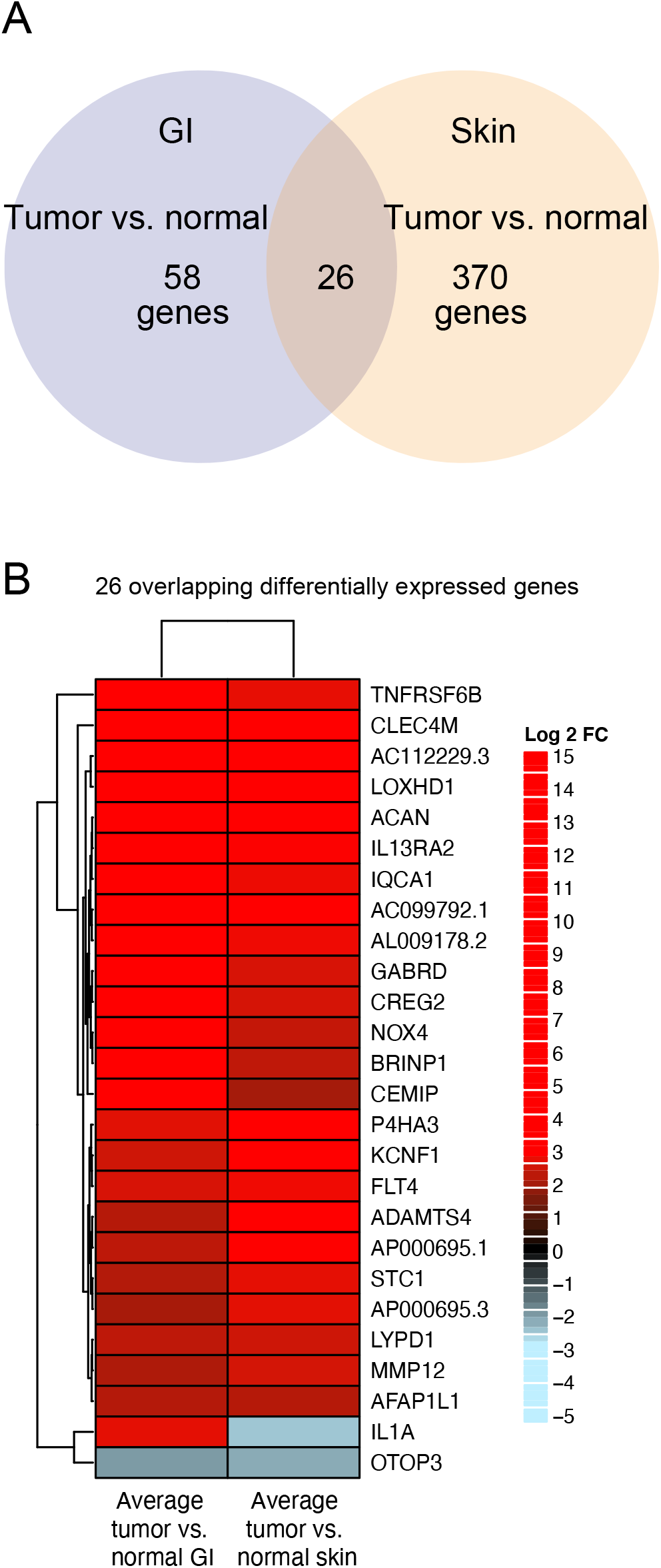
Differentially expressed genes from GI (KS tumor versus normal) samples and differentially expressed genes from skin (tumor vs normal in green) samples. The differential gene expression cutoff of log2 Fold Change > or < 2.0 and padj. < 0.05 between tumor vs normal in both GI and skin samples was used. A. Venn diagram shows unique and shared differentially expressed genes. B. Heatmap showing shared differentially expressed genes using average Log_2_ fold change values.

Inflammation plays a pivotal role in the pathogenesis of KS as well as other KSHV-associated diseases such as MCD, KICS and PEL. Inflammatory responses also play a role in the host response to KSHV, but the virus has developed several mechanisms to evade such responses(34). Genes associated with inflammatory responses were explored to study differential expression between KS lesions and normal tissue samples (Fig. S1 and S2). Overall, many of these inflammatory genes were differentially expressed in skin KS, but not in GI KS. As expected, housekeeping (*HPRT1*, *GAPDH*, *HMBS*) genes were not significantly altered in expression when comparing KS lesions to their normal matched tissue samples (Figure 3A). Serum IL-6 and IL-10 levels are often elevated in KS and are especially elevated patients with other KSHV-associated diseases that occur concurrently with KS (3, 6, 26). *IL6* and *IL10* RNA levels were increased in skin KS (log2FC=2.1, adj. p=0.001; log2FC=2.1, adj. p=0.0009, respectively), but not significantly altered in GI KS lesions as compared to matched normal tissues. Furthermore, increased expression of IL6 correlated with increased total KSHV gene expression in skin KS (R^2^=0.67, p=0.004) but not in GI KS (Fig 3B). We also examined angiogenesis-related genes, since aberrant angiogenesis and increased vascularity are prominent features of KS lesions irrespective of their location (16, 35). VEGF receptors 1, 2, and 3 are encoded by *FLT1*, *KDR*, and *FLT4*, respectively. All 3 VEGF receptors were upregulated in skin KS lesions and *FLT4* (*VEGFR3*) was increased in both skin and GI KS (log2FC=2.8, adj. p=0.00005; log2FC= 2.6, adj. p=0.001, respectively (Figure 3A)). Other factors and networks associated with KSHV-pathogenesis, including gene expression associated with cytokine dysregulation, antiviral response, and immune activation in KS lesions were also investigated.

**Figure 3.**
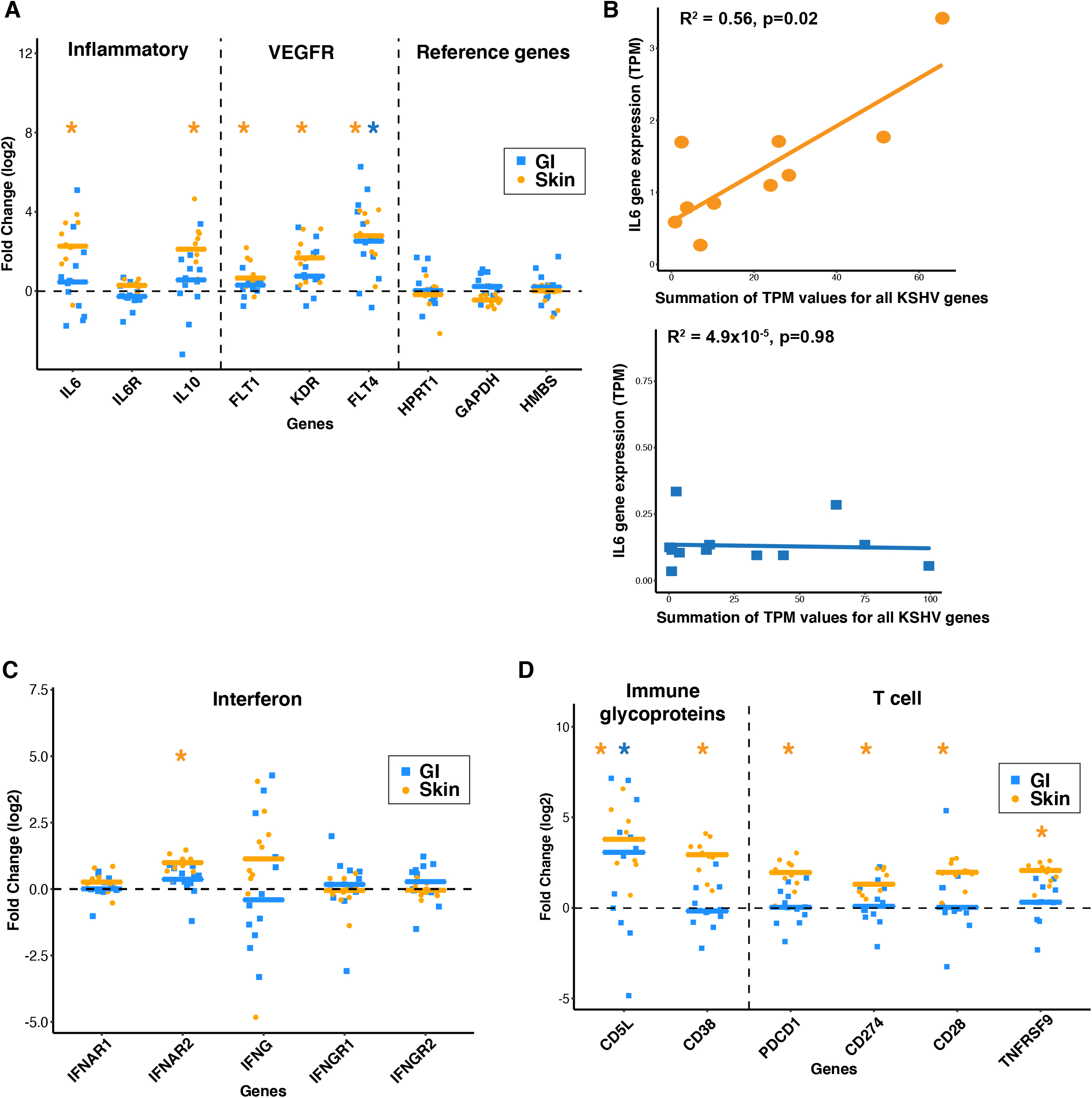
Selected genes associated with viral pathogenesis or immune responses are plotted. A, C, D. Asterisks (blue* for GI and orange * for skin) represent statistically significant (Student’s t test, p < 0.05) genes, in individual GI (tumor vs normal, log2 Fold Change, blue squares) and skin (tumor vs normal, log2 Fold Change, orange dots) samples. B. For only the KS tumor samples: transcripts per million (TPM) were collected for all KSHV genes on the horizontal axis and TPM for IL6 is shown on the vertical axis (orange for KS skin, blue for KS GI).

As a major antiviral host response, genes associated with the interferon (IFN) pathway were studied. Only *IFNAR2* showed consistent upregulation in skin KS samples (log2FC= 0.9, adj. p=0.00001, Figure 3C). There was a wide range of expression changes in interferongamma (IFN-γ), which has been implicated to be a factor in combating KSHV infection (36). In addition to interferon responses, the gene expression changes in genes involved in immune response were also evaluated. CD5L expression, which is predominantly released by macrophages, was increased in KS lesions in both skin and GI KS (log2FC=3.7, adj. p = 0.00008; log2FC=2.3, adj. p=0.04997, Figure 3C). CD38 is an active marker expressed in a variety of immune cells, including natural killer cells, B lymphocytes, and CD4+ and CD8+ T lymphocytes. There was a significant increase of *CD38* expression in skin KS (log2FC=2.8, adj. p=0.00002, Fig. 3D), but not of the GI tract. PD1 signaling can suppress T-cell responses. *PDCD1* (*PD1*) and *CD274* (*PD-L1*) were modestly increased only in skin KS lesions. CD28 is an important costimulatory molecule for survival and activation of T cells, which is regulated by TNFRSF9. Both of these genes followed the overall trend of increased expression in skin KS, but not in GI KS.

### Immune cell types within KS lesions and use of KSHV-infected lymphatic endothelial cell model

Given the heightened expression in markers associated with activated immune cells, we next applied the computational tool, CIBERSORTx(37), to interrogate changes in immune cell types in the KS lesions. This method analyzes the expression profile of specific marker genes from each cell type. We calculated the the relative amounts of cell type specific signatures KS lesions compared to their matched normal samples (Fig. 4A-B). Heterogeneity in the immune subtype signatures was noted across both skin and GI KS lesions. Follicular helper T cells and T regulatory cells were decreased in many GI samples, but this observation was not noted in the skin KS samples. There were elevated levels of resting memory CD4^+^ T cells in 5 of 12 GI KS samples as compared to the paired normal tissue and in 1 skin paired sample as compared to the paired normal tissue. Increased signatures of M1 macrophages were observed in the majority (7 of 10 samples, p=0.056) of skin KS samples.

**Figure 4.**
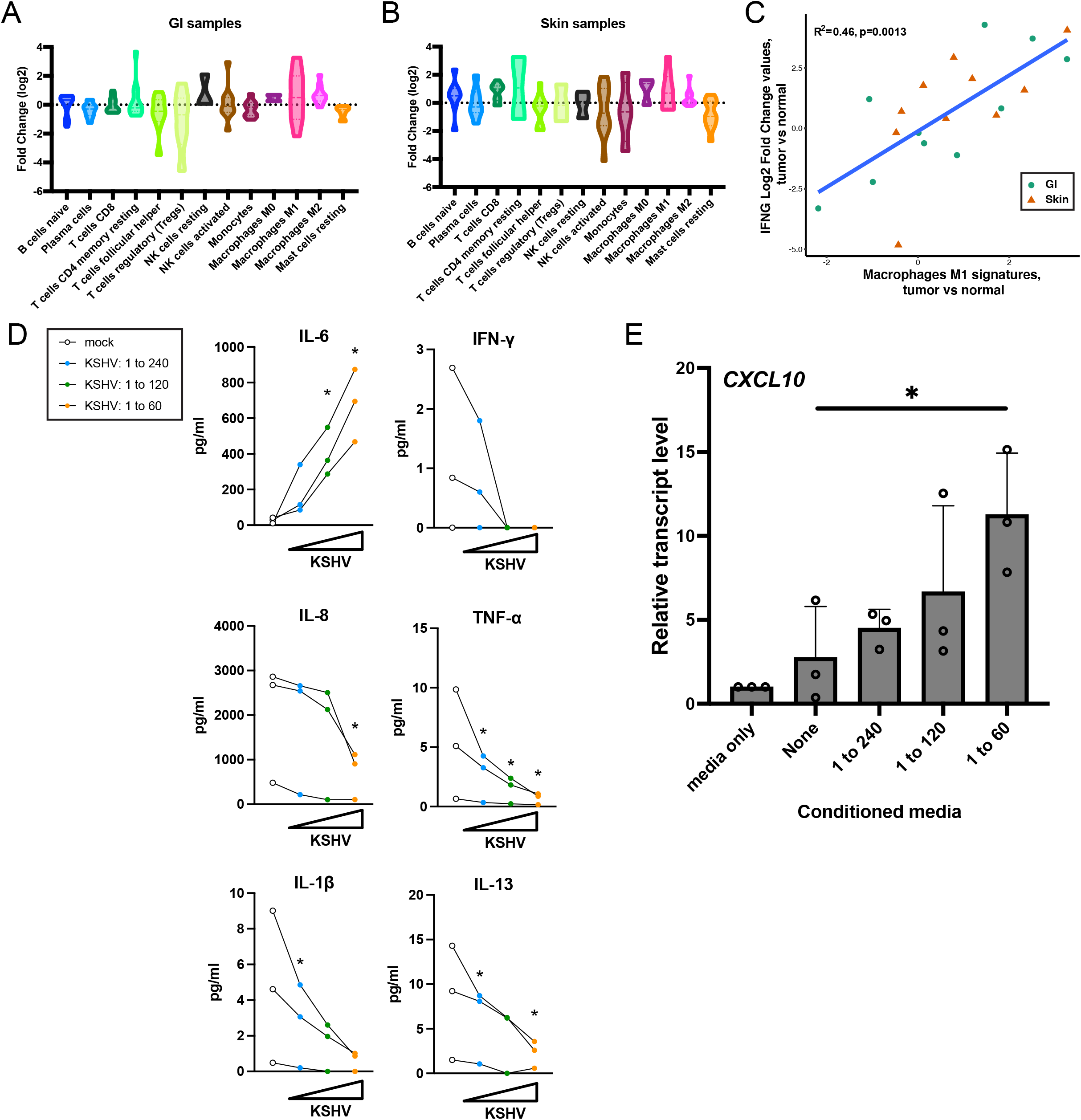
CIBERSORT analysis of KS lesions, cytokine secretion, and macrophage polarization. A-B. Fold changes of different immune cell types in tumor vs normal samples, were estimated by CIBERSORT. Violin plots show the increased or decreased fold change (log2) of each cell type signature of the KS lesions as compared to normal skin. C. Correlation analysis of changes in *IFNG* RNA expression (tumor vs. normal) and changes in M1 macrophage cell signatures (tumor vs. normal). D. Proinflammatory cytokine levels in conditioned media of KSHV-infected LECs after 3 days were quantitated with an electrochemiluminescence multiplex immunoassay. Each line represent a separate biological replicate. Only cytokines with concentration >5pg/ml in any of samples are shown. E. Macrophage polarization assays with conditioned media and a promonocytic cell line THP1. THP1 cells were treated with PMA and then incubated with conditioned media of KSHV-infected LECs (D) for 24 h. Transcript levels of M1 macrophage marker gene *CXCL10* was quantitated with RT-qPCR and normalized to *RPS13*.

Macrophages play key roles in antigen presentation, inflammation and phagocytosis. They exist in resting phases (M0) to polarizing states (M1 and M2) with varying phenotypes between M1 and M2. While M1, known as classical activated macrophages are pro-inflammatory and promote cell killing, M2 alternatively activated macrophages are anti-inflammatory and promote tissue healing. IFN-γ signaling promotes M0 macrophage polarization to M1 macrophages(38). Macrophage polarization is widely exploited by viruses through various mechanisms including manipulating cytokines (38). CIBERSORT analyses suggested that infiltrated macrophages in KS legions have different macrophage polarizations compared to normal samples. Within GI KS, RNA expression changes of IFN-γ varied widely as compared to their matched normal samples (Fig. 3C), and M1 macrophage signatures appeared variable in both tissues (Fig. 4A). Interestingly, there was a strong positive correlation between IFN-γ expression and M1 macrophage signatures in all tissues (p = 0.0013, Fig. 4C).

To further characterize the relationship between cytokines, macrophage polarization and the potential role of directing specific cell types in the KS tissue microenvironment, changes in cytokine levels following KSHV infection of endothelial cells were investigated. An initial question was whether IFN-γ expression levels are changed by KSHV infection of LECs or are the changing levels of IFN-γ expression due to other cell types (e.g. T and NK cells) in KS lesions. LECs were infected with KSHV in cell culture experiments and the conditioned media from these cells was collected. Changes in cytokine secretion from mock-infected and KSHV-infected LECs were measured at 3 and 5 days post-infection. Conditioned media of KSHV-infected LECs contained higher levels of IL-6 in virus dose-dependent manner (Fig. 4D). This increase in IL-6 levels in KSHV-infected endothelial cells was consistent with previous reports (39, 40). Other pro-inflammatory cytokines such as IL-8 and IL-1β were reduced in the conditioned media from KSHV-infected cells. Macrophage polarization to the anti-tumor M1 macrophage type is dependent on IFN-γ and TNF-α, and polarization to the pro-tumor M2 macrophage type depends on IL-4 and IL-13. Secretion of all these cytokines were reduced upon infection. IL-4 concentration was less than 5pg/ml in all samples and was not shown. To assess the effect of conditioned media from KSHV-infected endothelial cells on macrophage polarization, we differentiated a promonocytic cell line THP1 with conditioned media of infected LECs. THP1 cells can be differentiated to M1 or M2 types. Increasing the amount of KSHV in infections polarized THP1 differentiation more to a M1 profile as measured by an M1 macrophage marker gene, *CXCL10* (Fig. 4E). KSHV infection in LECs thus regulates secretion of pro-inflammatory cytokines and may promote M1 macrophage polarization. The M1 polarization observed in KS lesions was unlikely due to changes in IFN-γ secretion from KSHV-infected endothelial cells since IFN-γ secretion levels only decreased with KSHV infection in LECs (Fig. 4D). These cell culture experiments (Fig. 4D-E) showing a increase in an M1 macrophage marker was consistent with the increased M1 cell signatures observed in skin KS (Fig. 4B), compared to the matched normal skin tissue.

### KSHV gene expression in GI KS and skin KS lesions

KSHV gene expression patterns within the patient samples were also evaluated. In these bulk RNA sequencing analyses, these patterns represent an average expression profile of the mixture of all the KSHV-infected cells and this analysis cannot readily distinguish between low expression of a viral gene across many cells versus high expression of the same viral gene in a few cells in the lesion. KS is a heterogeneous tumor, previously noted to be largely comprised by cells expressing latent antigens with a subpopulation of cells expressing lytic antigens, as classified in tissue culture infection systems. We investigated whether these canonical latent and lytic gene expression patterns were observed in KS biopsies. Among GI KS and skin KS, there was heterogeneity with respect to the KSHV expression patterns in the heatmap analyses (Fig. 5A,B). In general, three patterns were detected across multiple lesions. First, there was expression of T1.1/PAN, a lytic marker, with little expression of other lytic genes. Second, expression of traditional latency genes (LANA, ORF72, K12, K13) were found, as expected. Third, there was moderate to high expression (TPM greater than 0.1) of more than 10 KSHV genes in 6 of 12 GI lesions and 5 of 10 skin lesions. The analyses of KS lesions identified some surprising trends in KSHV gene expression. First, high levels of ORF75 were observed when many other lytic genes were not found to be expressed. Second, the expression of ORF72/vCyclin D and LANA/ORF73 did not appear to correlate, despite being adjacent genes in the latency locus.

**Figure 5.**
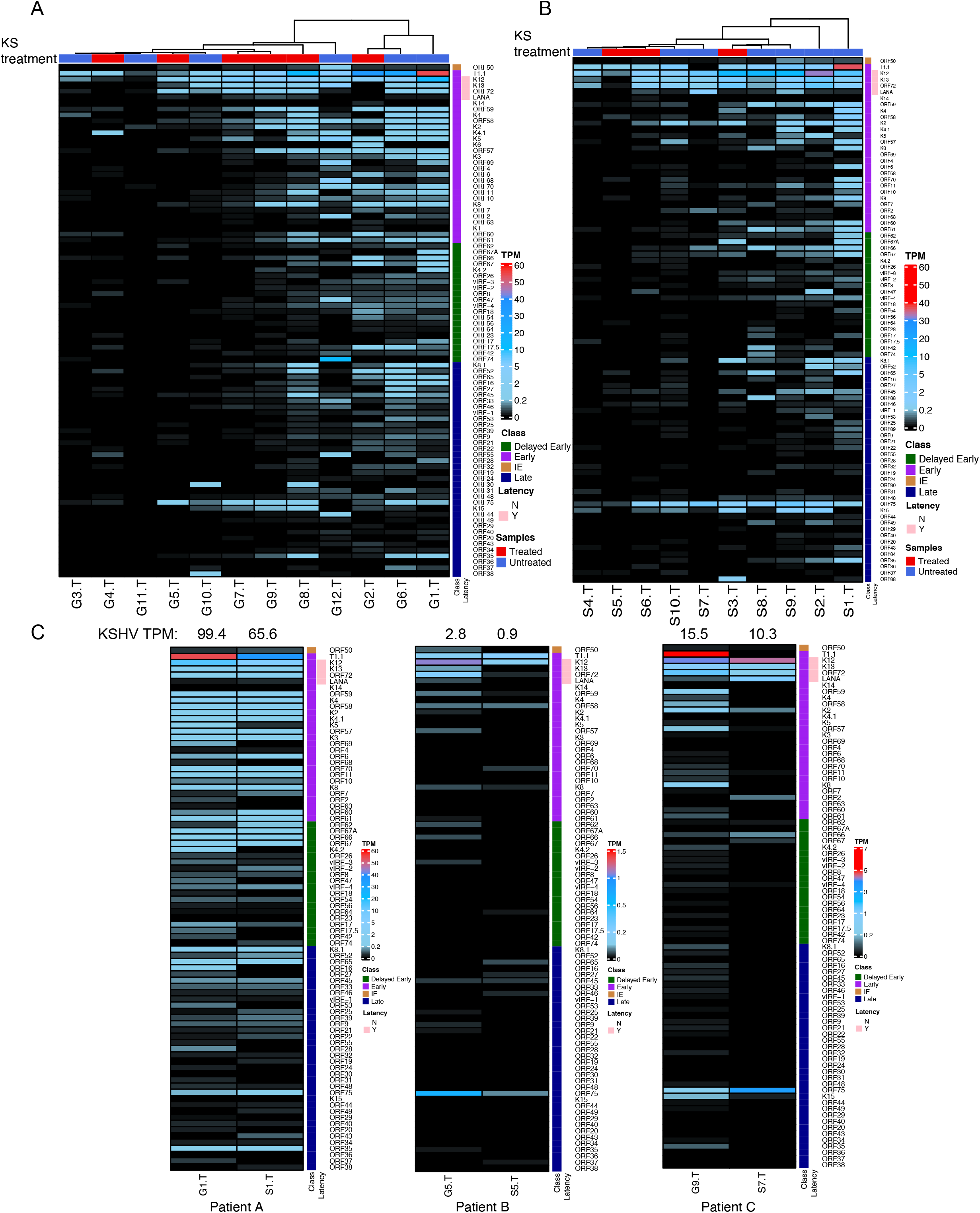
Heatmaps of KSHV gene expression KS tumors in GI (A) and skin (B) samples. Samples were either from participants untreated for KS (blue) or treated (red). Heatmap shows TPM expression from black (low) to medium (light blue) to high (red). ‘Class’ represent different stages of KSHV genes classification based on pervious studies as Immediate Early (brown),Early (purple), Delayed Early (green), and Late (blue). The latency genes were represented as not latent (N-white) and latent (Y-pink). C. KSHV gene expression heatmaps from the same matched participants. KSHV viral genes expression (Transcript per million (TPM) values (low-black, medium-blue, high-red) in GI and skin tumor samples from the same participants.

Three participants provided both GI and skin KS lesions at the same timepoint. We noted similarities in the expression of ORF75 and K12, irrespective of location in these participants (Fig. 5C). The underlying clinical characteristics may have had an impact in the overall KSHV gene expression in both the skin and GI KS lesions in these individuals. Though all 3 patients had uncontrolled HIV at the time of sample collection, with a CD4^+^ T cell count of <50 cells/μL, Patient A (Figure 5C – left) had no concurrent KSHV-associated diseases and had undetectable levels of circulating KSHV PBMC levels. This individual had uniform expression of lytic and latent KSHV genes in both samples. The remaining 2 patients had clinical and laboratory findings consistent with KICS and had greater KSHV gene expression in the GI samples. Among all 3 participants high levels of ORF75 were noted in both skin and GI samples, which is unusual as it has been described as a late gene and only expressed in lytically-infected cells. Interestingly, few other late genes were expressed when high ORF75 expression was found. In two of three participants, there was higher T1.1/PAN expression in the GI KS when compared to skin KS (Fig. 5C). Finally, in two of three participants, there was high expression of K2/viral IL6.

### KSHV gene expression in tissue sections

To determine whether ORF75, a marker of lytic replication was present in KS lesions, a randomly selected sample (noted in Figure 5 as S9T) was studied using immunohistochemistry and RNA *in situ* hybridization assays. Robust protein expression of KSHV LANA protein, and the endothelial cell marker CD31 were observed in sequential sections of the same area of tissue by in situ hybridization (Fig. 6A), findings consistent with KS pathology. Multiplex RNA *in situ* hybridizations were used to visualize viral transcript expression in a region verified to harbor KSHV infection based on LANA detection. KSHV ORF75 RNA was robustly detected in large clusters of cells that also expressed KSHV vIL6 RNA (Fig. 6B), in multiple areas of the biopsy. These data suggests that the high levels of transcripts detected by RNAseq are not limited to a small subpopulation of lytic cells and support the finding that ORF75 expression is widespread in KS skin lesions.

**Figure 6.**
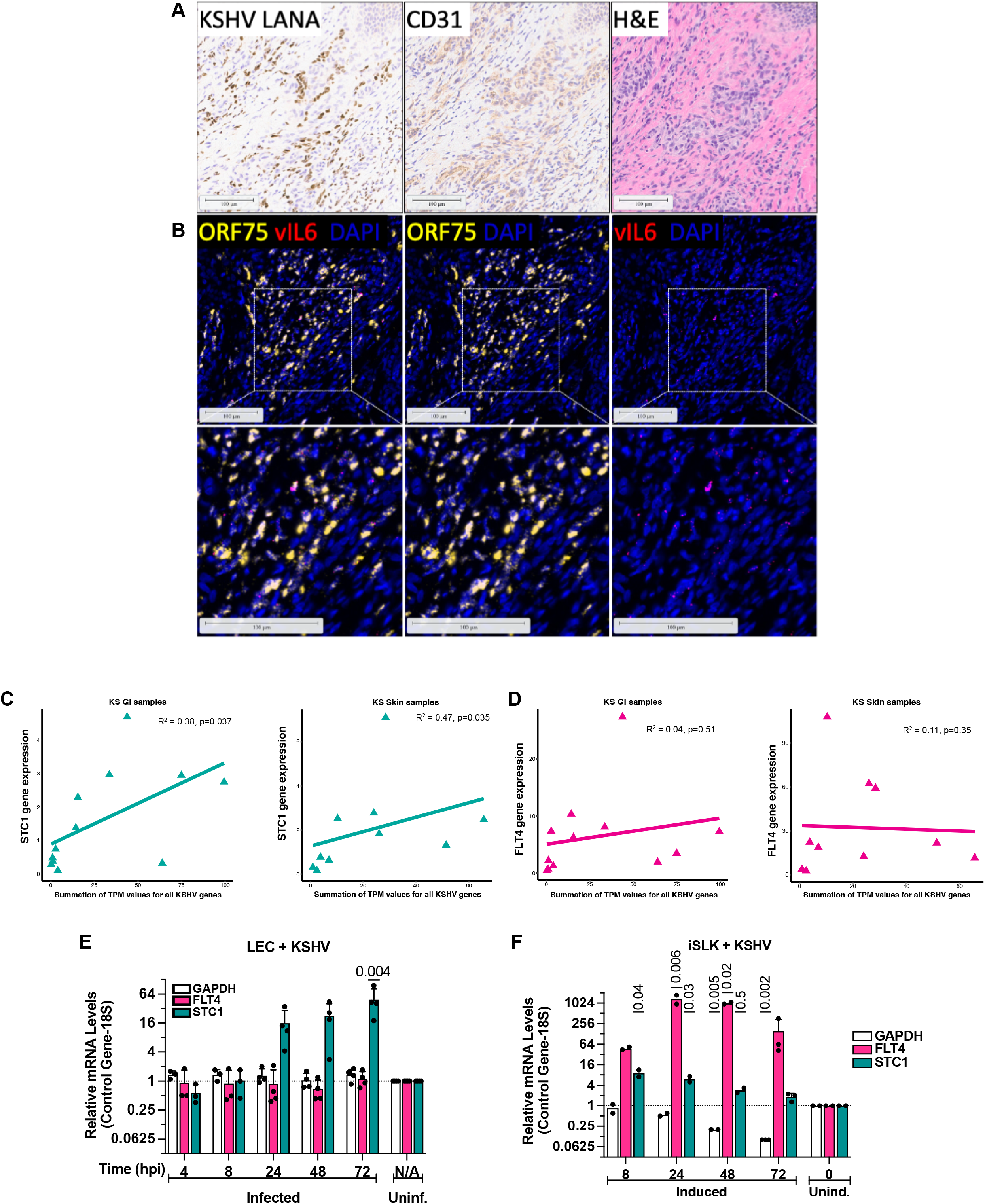
Expression of KSHV genes, *STC1*, and *FLT4* in KS lesions and laboratory infections. A-B. Sections from KS skin tumor sample S9T were used for protein and RNA expression detection assays. C-D. Total KSHV transcript per million (TPM) values versus STC1 and FLT4 TPM values were plotted. Spearman correlation analysis was performed for KS tumor samples. E. Human dermal lymphatic endothelial primary cells (HDLEC) were infected with KSHV and RNA expression was analyzed 4 to 72 hours post-infection using qPCR. Log2 fold changes were plotted comparing infected versus uninfected cells. F. KSHV-infected iSLK cells were lytically induced and RNA was harvested 8 to 72 hours after induction. RNA levels were measured using qPCR.

### STC1 and FLT4 expression in de novo infections

We sought to further investigate two cellular genes that were both upregulated in GI and skin KS, Stanniocalcin-1 (STC1) and Fms Related Receptor Tyrosine Kinase 4 (FLT-4) (Fig. 2B). First, STC1 was upregulated in both skin and GI KS (Fig. 2). Second, previous cell culture KSHV infections in primary endothelial cells demonstrated an increase in STC1 expression in KSHV-infected cells (41). Third, there was a correlation between total KSHV gene expression and STC1 expression in GI and skin KS (Fig. 6C). FLT-4 was also upregulated in GI and skin lesions (Fig. 2B). However, FLT4 expression did not correlate with total KSHV gene expression in KS lesions (Fig. 6D). This said, as a receptor for VEGF-C and VEGF-D, FLT-4 may be particularly important as a potential therapeutic target for KS it can be targeted by two FDA-approved medications (42).

The complexity of cell types and infections status in KS lesions led to the examination of changes in *STC1* and *FLT4* upon KSHV infection in more controlled laboratory experiments with LECs. Within this model, *STC1* expression strongly increased with *de novo* KSHV infection (Fig. 6E). In SLK cells, lytic induction of KSHV increased expression of *FLT4* around 1000-fold (Fig. 6F). Previously, KSHV infection has been shown to increase *FLT4/VEGFR3* expression in TIME cells (43). Taken together, the cell culture infections correlated with the observations in GI and skin KS that showed an increase in expression of *STC1* and *FLT4*.

### Effects of STC1 and FLT4 on phagocytosis, and angiogenesis in LECs

Small interfering RNAs (siRNAs) were used to assess the impact of *STC1* and *FLT4* expression on viral infection and cell behavior. Primary LECs were transfected with an siRNA control (siNTC) or siRNAs targeting *STC1* (Fig. S3). Initial assays demonstrated that repression of *STC1* did not inhibit early viral entry (Fig. 7A) when measuring a GFP reporter expressed by the KSHV virus or viral genomes per cell shortly after infection. Additionally, siRNAs knocking down *STC1* correlated with decreased production of viral particles (average relative decrease of 29%, p=0.0093) at 5 days post-infection wth the 1:120 dilution of virus (Fig. 7B). At 3-days postinfection, KSHV gene expression was decreased with the strongest reduction of *STC1* by siRNAs. Specifically, there was a reduction in expression of a KSHV latent gene, LANA, with lower virus input (1:240) (Fig. 7C). The strongest reduction with repression *STC1* was found for the lytic switch gene, RTA, with the 1:240 dilutions of the KSHV stock. In other words, knocking down *STC1* with siRNAs appeared to also decrease expression of some KSHV genes. A technical limitation of this assay included the transient nature of siRNA knock down assays and the robust increase in *STC1* expression shortly after KSHV infection in LECs (Fig. 6C).

**Figure 7.**
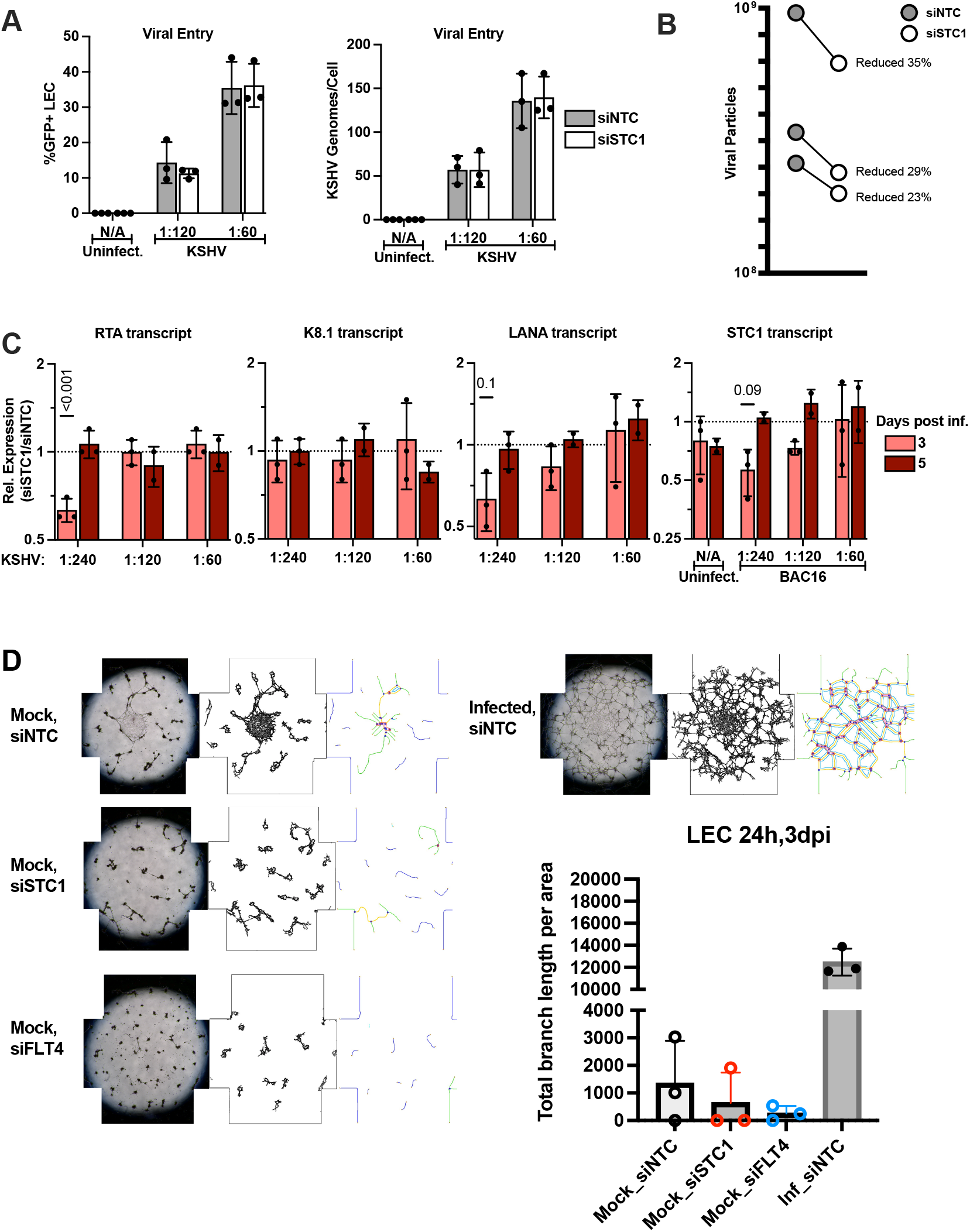
Repression of *STC1* and *FLT4* in primary human dermal lymphatic endothelial cells. A. Cells were transfected with siRNA non-targeting control (siNTC) or siRNAs targeting *STC1* (siSTC1). One day post-transfection, cells were infected with KSHV strain BAC16 (contains a GFP reporter). The percentage of GFP-positive cells was determined by flow cytometry (left). DNA harvested by cells was used in qPCR assays to measure viral genomes per cell at 1 day post-infection. B. New viral particles were measured from conditioned media from cells infected and transfected. Samples were collected 5 days after infection. Each line represents a separate biological replicate. C. RNA was purified from cells after transfection and infection and KSHV transcript levels were compared between siSTC1-transfected cells and siNTC cells. D. Endothelial cells were transfected once with siRNAs, infected, then seeded on Matrigel, and imaged by brightfield microscopy at 3 dpi. Tubule formation was assessed by image analysis software.

To determine whether STC1 and FLT4 play important roles in angiogenesis in LECs during KSHV infection, tubule formation assays were used and included computational analysis to measure the total length branched structures that were formed by LECs. As expected, the total branch length increased with KSHV infection at 3 dpi (Fig. 7D). Notably, siRNAs targeting either *STC1* or *FLT4* (Fig. S3) decreased the total branch length in uninfected LECs. Taken together, these results suggest that increased expression of *STC1* or *FLT4* in KS lesions may contribute to increased angiogenesis.

## Discussion

Kaposi sarcoma can manifest both in the skin and GI tract and have impact on morbidity and mortality of individuals in the United States, particularly among PLWH. These analyses investigated differences in GI and skin KS lesions to determine whether there are similarities or differences that would highlight pathways associated with KS pathogenesis and identify new KS therapeutic strategies. To our knowledge, this is the first study evaluating and comparing KS lesions from different disease sites. Additionally, this is the first study to investigate both cellular and KSHV transcripts from KS tissues within a well annotated cohort from the United States. There was considerable heterogeneity in the expression of host cellular genes between GI and skin lesions as compared to their matched normal tissues. Additionally, there was heterogeneity noted in immune profiles by KS site as well as KSHV-specific transcripts in both GI and skin lesions. There were 26 common cellular genes of interest identified between skin and GI lesions that implicate genes coding inflammatory cytokines as well as those associated with angiogenesis and immune regulation.

From our study, we note that there more complex patterns of KSHV gene expression in these KS lesions pushes beyond the traditional rigid definitions of latent versus lytic gene expression program. Detection of KSHV LANA protein is the standard assay for diagnosing KS. The expression analysis presented here suggests that if RNA expression techniques are utilized, additional KSHV genes may represent a more sensitive approach for detecting KS, potentially including ORF72 and ORF75. Though such technologies are limited by cost and analytical aspects in limited resource settings. There was higher expression of ORF72, compared to ORF73/LANA in both skin and GI KS lesions. This observation was consistent with a report from a previous study that analyzed KSHV RNA expression from KS tumors and suggests transcription initiation from an alternative promoter called LTd(33). The transcriptional regulation of promoter usage may differ between the KS lesions in this report and PEL cell lines. In the limited number of samples from skin and GI KS from the same patient, there were trends of more lytic expression in the GI KS lesions, compared to within skin KS lesions.

In this study, nineteen participants provided 22 samples and the median CD4^+^ T cell count of 38 cells/μL. Those who provided skin samples generally had well controlled HIV, whereas among participants who provided GI samples were noted to have uncontrolled HIV and all but one patient had concurrent KSHV-associated diseases, such as PEL, KICS or MCD. Therefore, the considerable heterogeneity between skin and GI samples may also be related to the underlying clinical characteristics and concurrent KSHV-associated process. In the presence of other KSHV-associated processes, elevated inflammatory cytokines (IL-1, IL-6 and IL-10) are notably high in the circulation(3, 26, 27). Skin KS lesions had higher IL-6 and IL-10 gene expression compared to normal tissue. In skin KS lesions, IL-6 gene expression had a positive correlation with the sum of KSHV transcripts. This was not the case in the comparison between IL-6 and KSHV genes in GI lesions. This differential correlation highlights the role of KSHV infection in regulating IL-6 expression specifically in skin KS lesions. In these analyses, transcripts with higher IL-1A levels were noted in GI KS lesions as compared to the normal mucosa, with lower levels in the skin KS lesions as compared to the corresponding normal tissue. However, this was not the case with IL-6 and its association with elevated expression levels, which were similar in both KS sites in comparison with respective normal samples.

With respect to other cellular genes, the gene expression profiles were in line with previous studies that have shown that pathways associated with angiogenesis are important in KS pathogenesis. Among genes notable in both GI and skin KS lesions as compared to their respective normal tissue, both FLT4 and STC1 were increased; this has also been noted in previous studies in HIV-associated KS(44, 45). FLT1 was also higher in skin tissue as compared to the normal skin. Studies investigating the role of both STC1 and FLT4 in KSHV infected LECs demonstrated that these two factors increased in KSHV infections. A previous study of 4 cisgender male participants with HIV and cutaneous KS identified cellular genes associated with lipid and glucose metabolism; however, this was not observed in the current study(45). Several interferon genes were studied. *IFNAR2* alone was elevated in the skin KS lesions as compared to the normal skin. A wide range of *IFNG* expression changes were observed in both skin and GI KS lesions as compared to the normal tissue. Among these tissues that were evaluated, it is interesting to note that among the KSHV genes, the viral interferon regulatory factors (vIRF) expression, which is known to regulate this antiviral response, was variable in its expression in all tissues(46, 47).

The differences in skin and GI lesions gene expression may be associated with the immune profiles in the respective tumor microenvironments. A recent study using single cell sequencing methods of KS among patients who were recently diagnosed with HIV and KS who were antiretroviral therapy naïve demonstrated an abundance of T cells and macrophages(48). A previous study of KS lesions of the skin of patients with endemic and epidemic KS from sub-Saharan Africa demonstrated the distribution of immune cells by infection of KSHV cells. In these analyses, we noted classically activated M1 macrophages were evenly distributed in KS lesions, whereas alternatively activated macrophages were notable in areas that did not have KSHV infected cells(49). M1 macrophages are often amplified by IFN-γ signalling to promote cell killing(50). This study also highlighted the abundance of CD8+ T cells in KS lesions compared to normal skin but this was not related to KSHV infected cells. In these analyses, there were differences noted in the skin and GI KS lesions compared to the normal counterparts with higher M2 macrophages and higher resting memory CD4^+^ T cells particularly in GI KS lesions. Though this was not specifically correlated with KSHV infection levels or spatial configuration, it suggests that these immune cells may play a role in regulating the immune environment. Moreover, these findings demonstrate that such differences may be related to site specific immune environments between the skin and GI tracts rather than associated with KSHV or HIV infection. In analyses of paired skin and GI samples from 3 participants show variation in the immune profiles and a few similarities in the KSHV gene expression between sites. One hypothesis was that increased IFN-γ expression in KS lesions might be increasing M1 polarization. However, IFN-γ secretion levels dropped with KSHV infection in our experiments using LECs. In our results using conditioned media from KSHV-infected LECs, we did find an increase in an M1 macrophage marker. Taken together, these results suggest that factors other than IFN-γ in the secretions from KSHV-infected cells may be promoting M1 polarization.

In exploring angiogenesis within LECs using tube formation assays, inhibition of STC1 and FLT-4 led to decrease in tube formation. These findings may impact potential therapeutic options for KS. The increased expression of genes associated with angiogenic pathways raise the possibility of exploring anti-angiogenic therapies. Bevacizumab (a monoclonal antibody against VEGF-A both alone (51) and in combination liposomal doxorubicin (52) led to some responses, but less than other approved therapies responses and challenging toxicity profile in patients with KS. There are newer drugs active against FLT4 or VEGFR3 that are licensed for use in medullary thyroid cancer(42). Other findings with possible therapeutic include targets for IL1A, such as anakinra, which has been shown to treat a wide range of inflammatory syndromes(53). This may be useful in cases where patients present with KS in the GI tract alone. Previous reports identified genes important for survival of KSHV-infected cells in B cells(54) and endothelial cells(55). Some of these genes were upregulated in these analyses of skin KS (*BTAF*, *VMP1*, *DCHS1*, *VCAM1*, *PLXND1*) (Fig. S4). It will be of interest to determine if inhibiting these genes in KS can promote cell death of KSHV-infected cells. Additionally, the immune profiles seen in these analyses reinforce the use of chemotherapy sparing options, such as immunotherapy, which may increase T cell stimulation and reduce markers of T cell exhaustion associated with chronic HIV and KSHV infection(56).

The heterogenous clinical characteristics, such as the presence of concurrent KSHV associated disease, KSHV levels and HIV characteristics, within these analyses is a limitation and this makes it difficult to make a general conclusion about KS lesions both of the skin and GI tract. The inclusion of three participants who have both GI and skin KS collected in tandem help control for some of these issues. These analyses did not explore the role of HIV coinfection in KS. However, with only 2 participants without HIV, it is challenging to make general statements on this issue and may be explored in future analyses. Despite these limitations, these analyses provide a broad overview for specific analyses in future studies, specifically for GI KS that is associated with severe manifestations of this disease and rarely studied.

In summary, this study evaluates a selection of participants from the United States presenting with cutaneous and visceral KS. These analyses highlight differences in the KS pathogenesis pathways and immune profile by site. Future studies include investigating the spatial association of these gene expression patterns to better understand how KSHV-infected cells can impact these changes.

## Methods

### Participants and clinical samples

Individuals with KS under the care of the HIV/AIDS Malignancy Branch at the National Cancer Institute were included. At the time of evaluation, participants had a 6mm punch biopsy of cutaneous KS over the lower limb, with normal appearing skin obtained within the same limb. If participants had gastrointestinal symptoms, an endoscopy or colonoscopy was performed and the operator visually identified KS lesions that were biopsied. An adjacent area of normal mucosa was obtained at the same time. All KS biopsies that were obtained for these analyses were divided and a portion was also sent to the laboratory of pathology for histological confirmation. All participants were consented to protocols for tissue procurement (NCT00006518) and genomic sequencing of KS and other KSHV-associated diseases (NCT03300830). Both protocols were approved by the NCI Institutional Review Board. All enrolled participants gave written informed consent in accordance with the Declaration of Helsinki.

Clinical and HIV characteristics were obtained at the time of biopsy collection. KSHV viral load (VL) in peripheral blood mononuclear cells (PBMCs) was assessed by quantitative real-time polymerase chain reaction as previously described (57).

### RNA-sequencing and analysis

Tissues were stored in RNAlater and lysed with Trizol. Tissues were homogenized and extracted for total RNA with Direct-Zol Miniprep kit (Zymo). Ribosomal RNA was removed and sequencing libraries were prepared using Illumina TruSeq Stranded / NEBnext Ultra Low Input Total RNA Library Prep and paired-end sequencing. The reads from the FASTQ files (reads are generated by Illumina HiSeq4000) were trimmed of Illumina adapters using *cutadapt* (1) with default parameter except for --q 10 and --minimum-length 25. We generated a ‘combined genome reference’ by catenating the human reference genome (GRCh38) and KSHV reference genome (NC009333). The reads (average 45.9 million reads per sample) were aligned to the combined genome reference using STAR (version 2.7.8a (2)). The transcriptome bam files created by STAR was used to estimate the read counts and Transcript per million (TPM) values at the gene level using RSEM v1.3.2 (3). The GENCODE v30 reference gene annotation (GTF) combined with the KSHV gene annotation were used for annotation using STAR and RSEM.

To find differentially expressed genes (DEGs) between tumor and normal samples, pairwise comparison was carried out using DESeq2 (4). DEGs were selected for the further study based on log2FoldChange > or < 2 and adjusted p value < 0.05. TPM values was used to generate scatterplot. All the plots were generated using R package ggplot2. The log2 foldchange values were considered for the jitterplot by comparing tumor vs normal using edgeR program (5). The TPM values of viral genes was considered to generate the heatmap using ComplexHeatmap package (6). CIBERSORTx software predictions were based on TPM values using the leukocyte signature matrix (LM22) (7).

### Human cytokine analysis

Conditioned media (100 μl) was used for measuring specific cytokines with V-PLEX Proinflammatory Panel 1 Human Kit (Meso Scale Diagnostics). Assays were performed by Clinical Support Laboratory (NCI-Frederick).

### Staining for proteins and RNA transcripts in tissue sections

Staining for LANA (Leica mouse monoclonal PA0050 ready-to-use) and CD31 (Abcam rabbit polyclonal ab32457;1:200 dilution) was performed on a fully automated BondMax autostainer (Leica) and detected with DAB using the HRP Polymer Refine Detection Kit (Leica) and hematoxylin counterstaining. FFPE sections of 5 μm were prepared from fixed tumor samples that were deparaffinized and rehydrated with a series of ethanol washes to deionized water. Sections were subjected to citrate (CD31) or EDTA-based (LANA) antigen retrieval for 20 minutes prior to immunostaining. Slides were incubated with primary antibody for 15 minutes (LANA) and 30 minutes (CD31). All slides were scanned and analyzed on an Aperio ImageScope scanner (Leica) and reviewed by pathologist L. Bassel. Human herpesvirus 8 ORF75 and viral IL-6 expression was detected by staining 5 um FFPE tissue sections with RNAscope 2.5 LS Probes V-HHV8-ORF75 (ACD, Cat# 562058) and V-HHV8-K2-C3 (ACD, Cat# 897938-C3) using the RNAscope^®^ LS Multiplex Fluorescent Assay (ACD, Cat# 322800) with a Bond RX auto-stainer (Leica Biosystems) with a tissue pretreatment of 15 minutes at 90°C with Bond Epitope Retrieval Solution 2 (Leica Biosystems), 15 minutes of Protease III (ACD) at 40°C, and 1:750 dilution of TSA Plus-Cyanine 3 (AKOYA Biosciences, cat# NEL744001KT) and TSA Plus-Cyanine 5 (AKOYA Biosciences, cat# NEL745001KT), respectively. The RNAscope^®^ 3-plex LS Multiplex Negative Control Probe (Bacillus subtilis dihydrodipicolinate reductase (dapB) gene in channels C1, C2, and C3, Cat# 320878) was used as a negative control. The RNAscope^®^ LS 2.5 3-plex Positive Control Probe-Hs was used as a technical control to ensure the RNA quality of tissue sections was suitable for staining. Slides were digitally imaged using an Aperio ScanScope FL Scanner (Leica Biosystems).

### Cell culture, reagents, nucleofection, and KSHV infection

Human dermal lymphatic endothelial cells (HDLECs) were obtained from Promocell and passaged in EGM2 medium (Lonza) for up to 5 passages, with passages 3 to 5 used for experiments. ON-TARGETplus nontargeting control siRNA and ON-TARGETplus SMARTpool siRNA targeting *STC1* and *FLT4* were obtained from Dharmacon/Horizon Discovery. HDLEC *de novo* infections were carried out using KSHV-BAC16 (concentrated by ultracentrifugation), diluted in EGM2 medium. Polybrene was added at 8 μg/ml. Uninfected samples were included as negative controls. After 5-6 hours of incubation, cells were washed and overlaid fresh media. RNA was harvested and extracted using Direct-zol kit (Zymo) with DNAse treatment. Viral entry was measured by detecting GFP-positive cells using flow cytometry. Viral entry was also measured by treating cells with Proteinase K, then phenol-chloroform-isoamyl alcohol, and ethanol precipitation of DNA. Viral DNA was measured by qPCR. Viral particles were collected from conditioned media after filtering with 0.45 μm filter. Filtered material was digested with DNAse I and virion DNA was purified using Proteinase K, then phenol-chloroform-isoamyl alcohol, and ethanol precipitation of DNA.

### Quantitation of mRNAs and proteins

Quantitative reverse transcription-PCR (RT-qPCR) was performed using 200-500 ng RNA and random primers with an Applied Biosystems high-capacity cDNA reverse transcription kit. SYBR green assays (FastStart universal SYBR green master mix; Roche or Thunderbird SYBR qPCR master mix, Toyobo) and TaqMan assays (TaqMan Universal PCR master mix, no AmpErase UNG; Applied Biosystems) were performed using the ABI StepOnePlus real-time PCR system (Applied Biosystems). Relative mRNA levels were computed using the threshold cycle (ΔΔCt) method with *RPS13* as a reference gene.

### Tube Formation Assays

HDLEC cells were transfected with siRNAs, then infected with KSHV BAC16, then plated on Cultrex Ultimatrix RGF BME for tube formation assays. Cells were stained with Calcein AM and microscope images were analyzed by ImageJ and Angioanalyzer(58).

## Statistical analyses

For genomic-level expression analysis, adjusted p-values were calculated using the Benjamini and Hochberg method. For smaller sets of comparisons, Student’s *t* tests was used. To determine the correlation between human and KSHV viral gene expression, Spearman correlation analysis was used.

## Acknowledgements

We would like to thank Amanda Day and Wendi Custer Lawrence for their technical assistance.

## Notes

### Competing Interest Statement

The authors have declared no competing interest.

