## Supplementary material for "Transcriptional analysis identifies overlapping and tissue-distinct profiles between Kaposi sarcoma tumors of the skin and gastrointestinal tract": Table S1

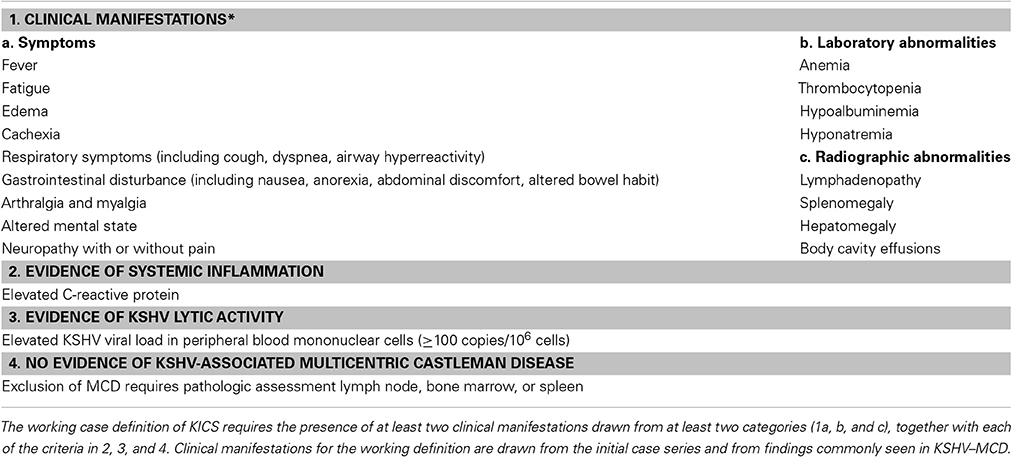
**Supplementary Table 1:** KSHV inflammatory cytokine syndrome case definition from Polizzotto MN, Uldrick TS, Wyvill KM, Aleman K, Marshall V, Wang V, et al. Clinical Features and Outcomes of Patients With Symptomatic Kaposi Sarcoma Herpesvirus (KSHV)-associated Inflammation: Prospective Characterization of KSHV Inflammatory Cytokine Syndrome (KICS). Clin Infect Dis. 2016;62(6):730-8.
